# Tryptophan recovers sensitivity to cell membrane stress in *Saccharomyces cerevisiae*

**DOI:** 10.1101/344366

**Authors:** Lea Schroeder, Pauletta Lazarevskiy, Amy E. Ikui

## Abstract

Sodium dodecyl sulfate is a detergent that disrupts cell membranes, activates cell wall integrity signaling and restricts cell growth in *Saccharomyces cerevisiae*. However, the underlying mechanism of how sodium dodecyl sulfate inhibits cell growth is not fully understood. Because deletion of the *MCK1* gene leads to sensitivity to sodium dodecyl sulfate, we implemented a suppressor gene screening revealing that the *TAT2* tryptophan permease rescues cell growth to sodium dodecyl sulfate-treated *Δmck1* cells. Therefore, we questioned the involvement of tryptophan in the response to sodium dodecyl sulfate treatment. In this work, we show that *Δtrp1* cells have a disadvantage in the response to sodium dodecyl sulfate compared to auxotrophy for adenine, histidine, leucine or uracil. While also critical in the response to tea tree oil, *TRP1* does not avert growth inhibition due to other cell wall/membrane perturbations that activate cell wall integrity signaling such as calcofluor white, Congo Red or heat stress. This implicates a distinction from the cell wall integrity pathway and suggests specificity to membrane stress as opposed to cell wall stress. We discover that tyrosine biosynthesis is also essential upon sodium dodecyl sulfate perturbation whereas phenylalanine biosynthesis appears dispensable. Finally, we observe enhanced tryptophan import within minutes upon exposure to sodium dodecyl sulfate indicating that these cells are not starved for tryptophan. In summary, our results expose a functional link between internal tryptophan levels and tryptophan biosynthesis in the response to plasma membrane damage.

## Introduction

In the wild, yeast experience a variety of external conditions that cause stress, such as changes in resource availability, temperature, osmotic fluctuations, oxidation, noxious chemicals, pressure and physical stress. The yeast cell wall and plasma membrane are the first defensive structures against external stress and are essential to acclimate to these conditions. In general, any perturbation that disrupts the cell wall or membrane function activates a multifactorial stress response in *Saccharomyces cerevisiae* (1, 2).

Sodium Dodecyl Sulfate (SDS) is a common household detergent that permeates cell membranes (3, 4), activates a stress response including Cell Wall Integrity (CWI) signaling and restricts cell growth (5). The CWI pathway is a kinase cascade that responds to cell wall/membrane perturbations in order to maintain cell integrity in yeast (1, 2). Treatments that damage the yeast cell wall or membrane such as chemicals like SDS (5), Calcofluor White (CFW) (6), Congo Red (CR) (7) and Tea Tree Oil (TTO) (8) or by growth at elevated temperatures (9), trigger the CWI pathway.

*MCK1*, the yeast homologue of the mammalian Glycogen Synthase Kinase-3 (GSK-3) (10, 11), is involved in a variety of stress response activities. Mck1p maintains genome integrity in response to DNA damage (12, 13) and is involved in the transcriptional regulation of stress response genes (14, 15). In addition, Mck1p is a downstream effector of CWI signaling activated by high temperature, osmotic stress or calcium stress (16, 17). Deletion of *MCK1* causes hypersensitivity to SDS (14, 16). We previously found that SDS induces cell cycle arrest during G1 phase via Mck1p (14). In order to understand the mechanism of cell growth inhibition by SDS, we implemented a suppressor gene screening using *Δmck1* cells in the presence of SDS. The screen revealed that the *TAT2* tryptophan permease rescued cell growth to SDS-treated *Δmck1* cells.

The high affinity tryptophan permease, Tat2p (Tryptophan Amino acid Transporter), is a constitutive permease regulated by the concentration of tryptophan in the media (18). The appropriate function and localization of Tat2p and/or the ability to biosynthesize tryptophan is required for yeast to survive a variety of stresses. In particular, perturbations that affect membrane stability have strong auxotrophic requirements for tryptophan. Yeast cells experiencing high pressure (19), a deficiency in ergosterol (yeast version of cholesterol) (20-22), organic acid stress (23) or ethanol stress, endure alterations to their membranes (24-27). In each case, tryptophan prototrophy or *TAT2* overexpression was a requirement for cell growth, indicating that tryptophan itself exhibits protection from membrane disruptions. In addition to these cell wall/membrane related stresses, it has been suggested that internal tryptophan levels influence growth recovery post DNA damage (28, 29).

Because our suppressor gene screening revealed *TAT2* and that tryptophan is linked to stress tolerance, we questioned the involvement of tryptophan in the recovery of cell growth in the presence of SDS. In this work, we show that SDS-induced growth inhibition can be overcome with exogenous tryptophan or tryptophan prototrophy. We found that tryptophan prototrophy exhibits protection from growth inhibition due to particular cell wall/membrane damaging agents that activate the CWI pathway, but not all treatments, suggesting that the need for tryptophan is autonomous from CWI activity. In addition to tryptophan biosynthesis, we show that tyrosine biosynthesis is also necessary for tolerance to SDS stress. Additionally, we determine that tryptophan import is not disrupted by SDS exposure but enhanced. These results provide an unknown connection to tryptophan and tyrosine in the protection from plasma membrane damage that is not due to general nutrient starvation and is independent of CWI signaling.

## Results and discussion

### Tryptophan availability recovers sensitivity to SDS

To affirm the rescue of *Δmck1* sensitivity to SDS with *TAT2*, we cloned *TAT2* into a pRS425 high copy plasmid and asked if *TAT2* alone rescues SDS-induced cell growth inhibition of *Δmck1* cells. Indeed, we found that the *TAT2* expressing plasmid conferred rescue to both *Δmck1* and *MCK1* cells in the presence of SDS (Fig 1A). It is known that during times of stress or nutrient starvation, Tat2p is sorted to the vacuole for degradation and then tryptophan uptake is maintained by Gap1p, the General Amino acid Permease (30-32). This may explain why the SDS-induced growth inhibition in *Δmck1* cells was only mildly rescued by *TAT2* overexpression.

**Fig 1.**
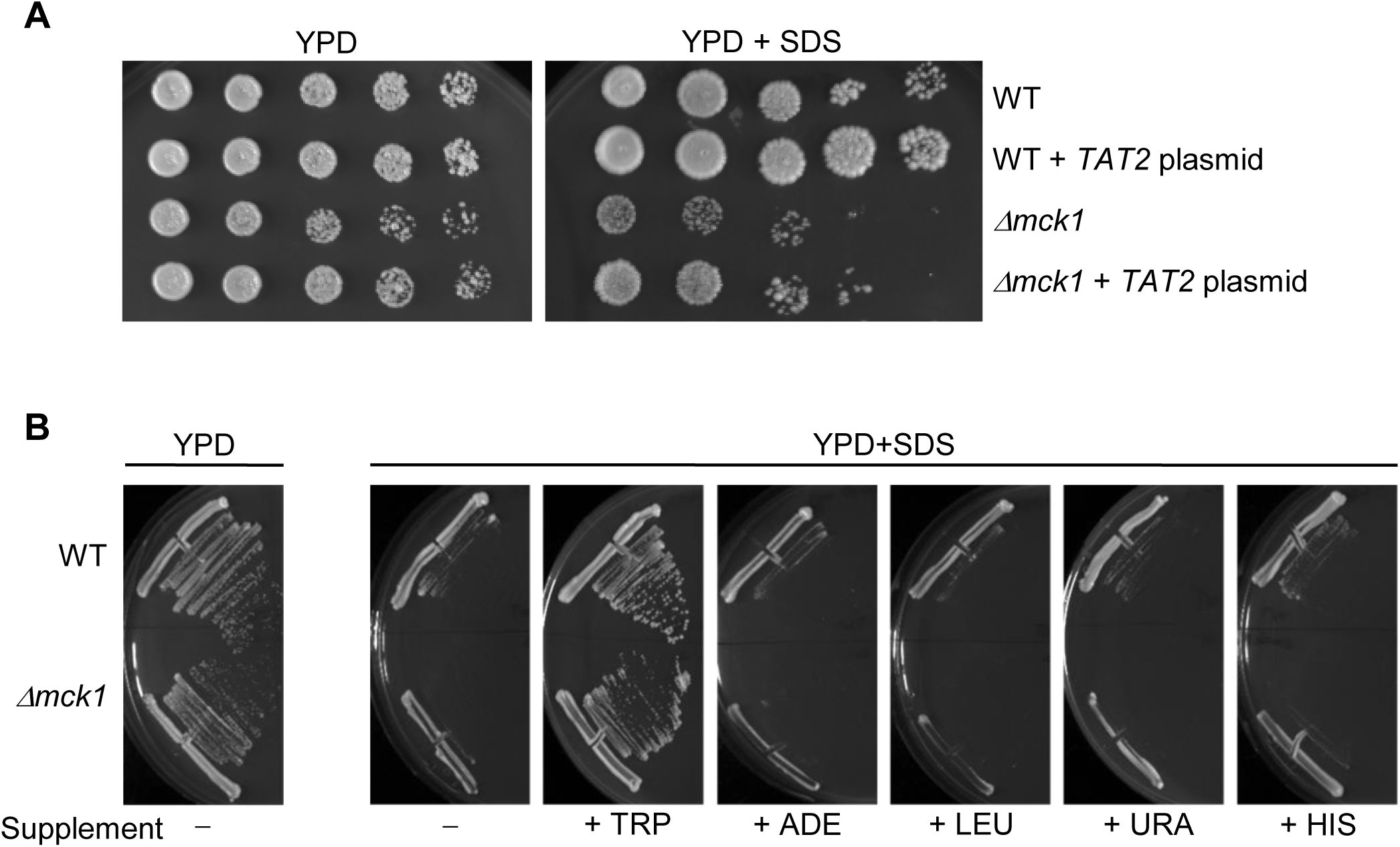
*TAT2*/Tryptophan rescues SDS sensitivity in Δ*mck1* cells. (A) Wild-type (WT) (*his3-11,15 leu2-3,112 trp1-1 ura3-1 can1-100*) and isogenic *Δmck1* cells transformed with *TAT2*/pRS425 plasmid were 10-fold serially diluted onto YPD or YPD containing 0.0075% SDS and incubated at 30°C for 2 days. (B) Wild-type (WT) (*his3-11,15 leu2-3,112 trp1-1 ura3-1 can1-100*) and isogenic *Δmck1* cells were streaked on YPD or YPD containing 0.0075% SDS plates supplemented with excess tryptophan, adenine, leucine, uracil or histidine. Plates were incubated at 30°C for 2 days.

To support the idea that cell growth sensitivity to SDS recovered by *TAT2* overexpression is due to tryptophan availability, we also observed that exogenous tryptophan recovered growth of both *Δmck1* and *MCK1* cells when we supplemented YPD plates containing SDS with excess tryptophan (Fig 1B). However, cell growth was still inhibited by SDS with the addition of exogenous adenine, histidine, leucine or uracil suggesting that recovery of SDS-induced growth inhibition is specific to tryptophan.

W303 and BY4741 are two commonly used lab strains of *S. cerevisiae*. It is already known that BY4741 cells that are auxotrophic for tryptophan are sensitive to SDS-induced cell membrane stress in liquid culture (33). We expanded these results by comparing BY4741 cells containing or lacking functional *TRP1* in a plate streaking assay. The presence of wild type *TRP1* also confersed resistance to SDS-induced growth inhibition on YPD plates containing SDS (Fig 2A, section 1 and 4). Considering this result and that *TAT2* overexpression rescued cell growth sensitivity to SDS in the W303 strain; we tested if SDS-dependent growth inhibition is a result of tryptophan auxotrophy alone. For this purpose, we used W303 cells that are prototrophic for the various markers *ADE2, HIS3*, *LEU2*, *TRP1* and *URA3* and compared the cell growth with isogenic tryptophan mutant cells, *trp1-1*, on SDS containing YPD plates. Indeed, the *trp1-1* mutation itself caused sensitivity to SDS in W303 cells (Fig 2A, section 5 and 6). From these results, we conclude that tryptophan auxotrophy alone was sufficient to cause sensitivity to SDS, indicating the significance of *TRP1* prototrophy for SDS resistance in multiple yeast backgrounds.

**Fig 2.**
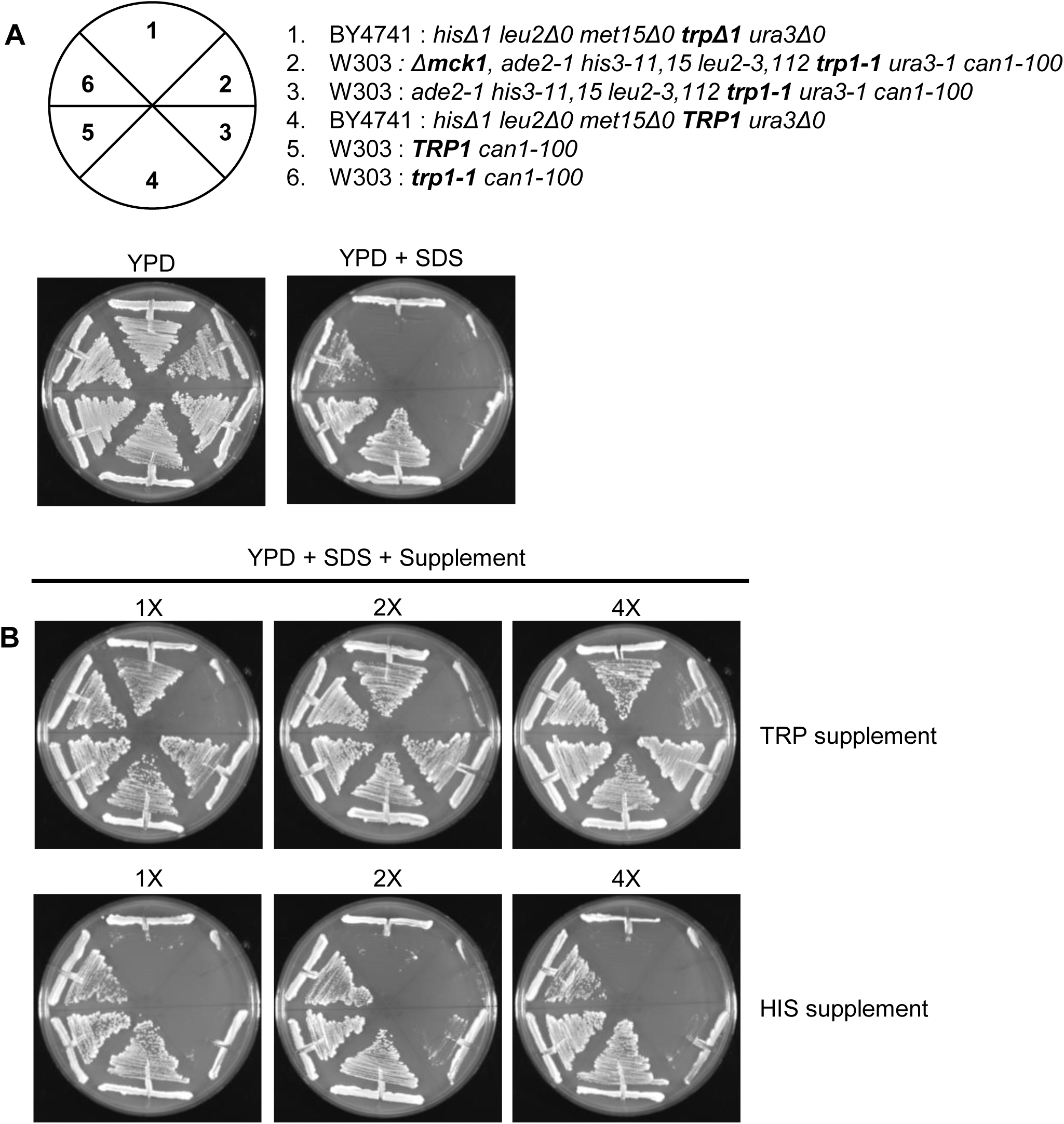
Tryptophan availability recovers growth sensitivity to SDS. (A) A schematic representing the location and cell genotypes used in the following experiments. The indicated cells were struck out on YPD or YPD containing 0.0075% SDS and (B) supplemented with increasing amounts of tryptophan or histidine (1X, 2X or 4X). The plates were incubated at 30°C for 2 days.

In addition, we tested if exogenous tryptophan rescues cell growth sensitivity to SDS in a dose dependent manner by comparing the same strains struck out on YPD plus SDS plates supplemented with increasing amounts of tryptophan or histidine. The growth of BY4741 and W303 cells disrupted for *TRP1* was uninhibited when excess tryptophan was externally available although growth was not dose dependent (Fig 2B, top row sections 1, 3 and 6). Histidine supplement was used as a control because it is an aromatic amino acid like tryptophan however it is not taken up through the Tat2p permease (18). Our results show that SDS-induced growth inhibition of W303 *trp1-1* cells was slightly alleviated by externally available histidine suggesting that a full complement of biosynthetic ability is favorable for the response to SDS (Fig 2B, bottom row section 6). The histidine supplement however, unlike the tryptophan supplement, did not suppress growth inhibition to either BY4741 or W303 cells harboring several auxotrophic markers (Fig 2B, bottom row sections 1 and 3) indicating that tryptophan is more important than histidine for the SDS response. These data show that cells harboring a nonfunctional *TRP1* are able to grow in the presence of SDS if a sufficient amount of tryptophan is externally available and that growth recovery is independent of strain background. *Δmck1* cells were also used as a control in these experiments and as expected, growth of the *Δmck1* cells was inhibited in the presence of SDS (Fig 2B, section 2). With excess tryptophan available, and not histidine, *Δmck1* cells showed mild growth recovery and *MCK1* cells that showed robust growth on YPD plates containing SDS (Fig 2B, section 2 and 3). This confirms our previous conclusion that *MCK1* contributes to the stress response in the presence of SDS in a way that might be independent from tryptophan availability.

### CWI activity is independent of tryptophan synthesis

These results prompted us to test further the prototrophic requirements for SDS resistance. W303 cells that are auxotrophic for all of the markers, *ade2-1, his3-11,15, leu2-3,112, trp1-1* and *ura3-1*, show growth inhibition on YPD plus SDS plates (Fig 3, row 4). The matched cells harboring a wild type copy of *TRP1* and are auxotrophic for the other four genes, showed robust growth in the presence of SDS whereas cells that were prototrophic for any other single mutant indicated besides tryptophan show growth inhibition in the same conditions (Fig 3, rows 1, 5, 6, 7 and 8). As an additional control, we used BY4741 cells that are prototrophic for *TRP1* and as expected these cells grow robustly at this concentration of SDS (Fig 3, row 2). These findings support the evidence that *trp1* auxotrophy has a more adverse effect to cells compromised with SDS than auxotrophy for adenine, histidine, leucine or uracil.

**Fig 3.**
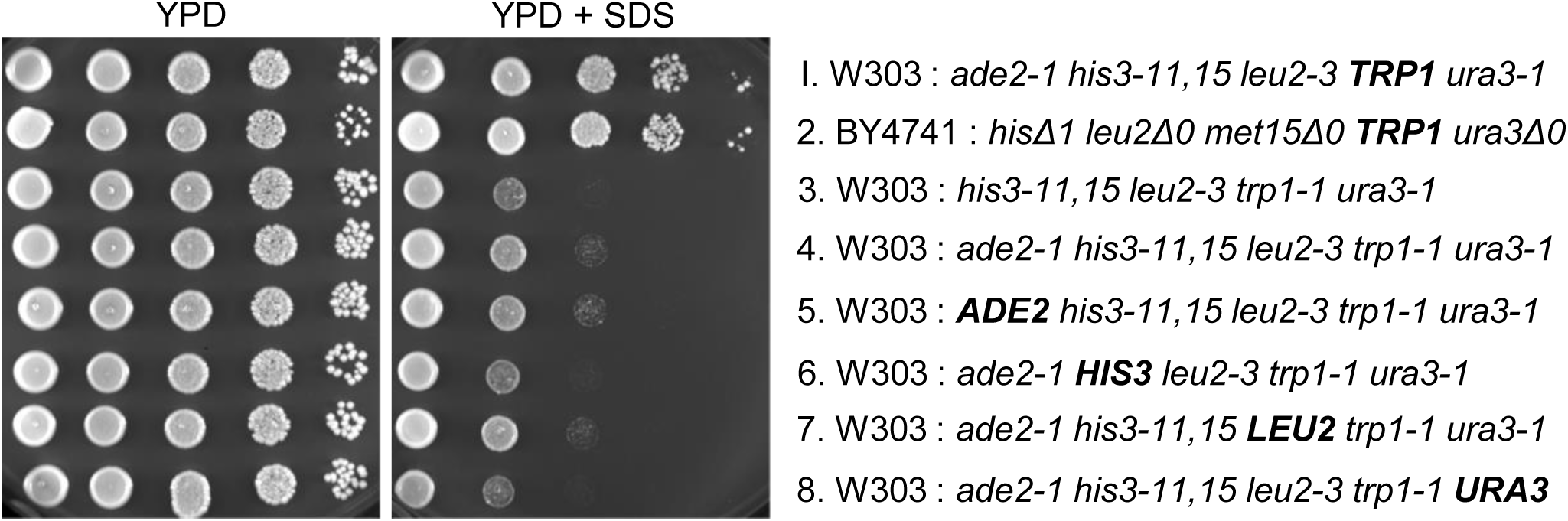
*TRP1* prototrophy confers growth advantage in the presence of SDS. The indicated cells were 10-fold serially diluted onto YPD or YPD containing 0.0075% SDS and incubated at 30°C for 2 days.

This result was also reflected in cells that are prototrophic for *ADE2, HIS3, LEU2, TRP1* and *URA3* (Fig 4A, rows 4, 5 and 6). The cells containing only a *trp1-1* mutation, were mildly sick in the presence of SDS whereas matched counterpart cells containing only a *his3-11,15* mutation did not show SDS-induced growth inhibition (Fig 4A, rows 4, 5 and 6). The increased resistance shown by the *trp1-1* cells compared to cells harboring multiple auxotrophies again suggests that prototrophy for all of these nutrients together are favorable in the response to SDS (Fig 4A, rows 1, 2, 3 and 8). However, *trp1-1* cell growth is still inhibited by SDS treatment compared to *his3-11,15* cells, providing further indication of the significance of *TRP1* prototrophy for tolerance to SDS.

Since SDS treatment activates the CWI pathway (34, 35), we wanted to know if tryptophan prototrophy can recover growth inhibition due to other activators of CWI signaling. CFW is a dye that interferes with cell wall assembly by blocking chitin polymerization, resulting in weakened cell walls (34, 36, 37). CR is a dye that interferes with cell wall assembly by binding to chitin and cellulose with high affinity (38, 39). Heat stress causes fluidization of the cell membrane and weakens the cell wall (40-42). TTO, an extract from the leaves of *Melaleuca alternifoli*, is a fungicide that disrupts cell membranes and mitochondrial functions (43, 44). Treatment of yeast with either dye, heat stress or TTO triggers the CWI pathway (8, 9, 40, 45).

We asked whether tryptophan prototrophy could recover growth sensitivity to cells challenged with 10ug/ml CFW or 10ug/ml CR. In contrast to SDS treatment, we found that the different varieties of W303 cells were as sensitive as each other upon CFW or CR treatment and this result is regardless of tryptophan prototrophy (Fig 4B). We also found that BY4741 cells are not as sensitive to CFW and CR as W303 cells (Fig 4B, row 7).

**Fig 4.**
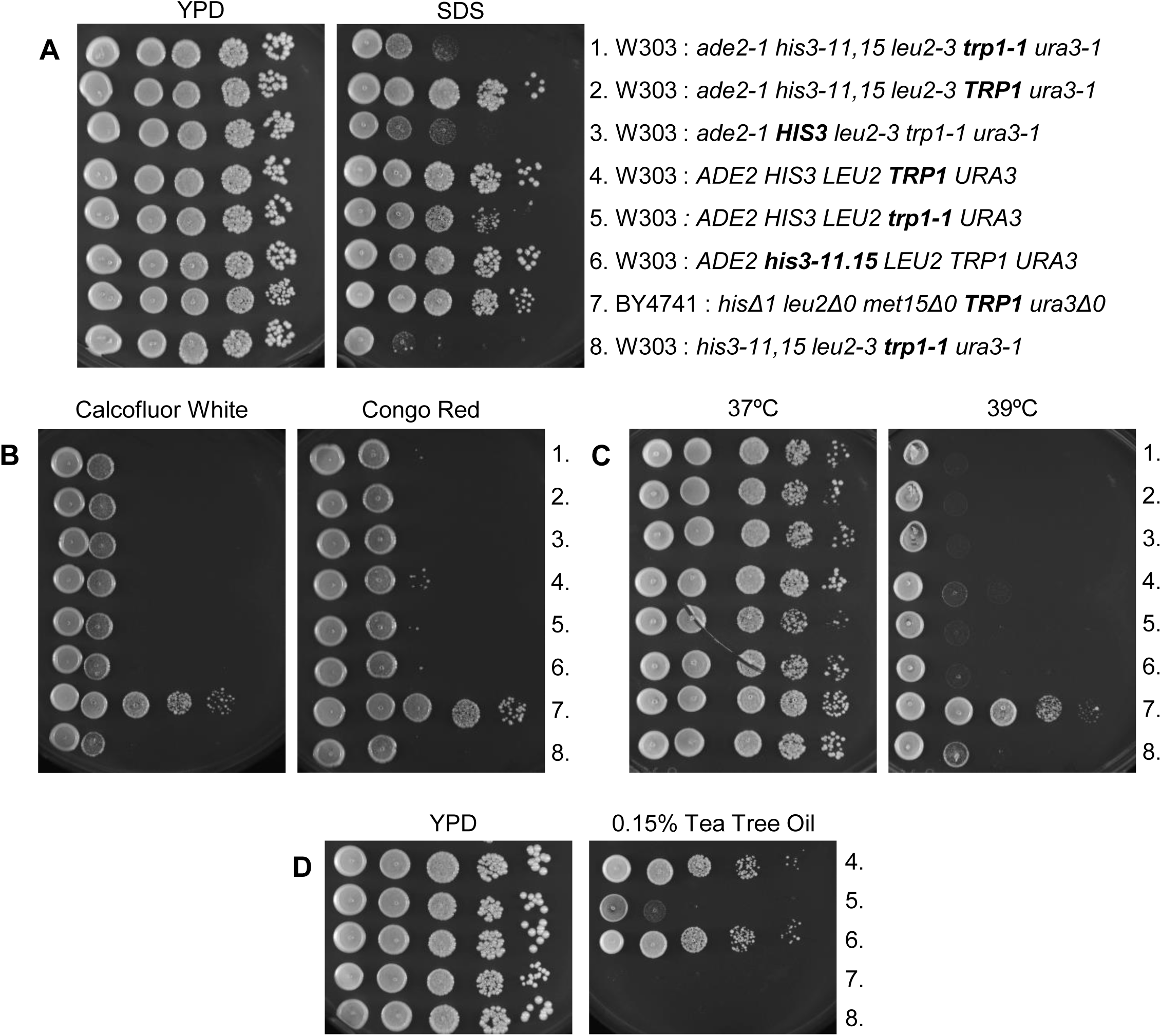
*TRP1* prototrophy recovers growth sensitivity due to some cell wall/membrane damaging treatments but not all. The indicated yeast cells were 10-fold serially diluted onto (A) YPD or YPD containing 0.0075% SDS (B) YPD containing 10ug/ml Calcofluor White or 10ug/ml Congo Red (C) YPD and incubated at 37°C or 39°C or (D) YPD containing 0.15% Tea Tree Oil. All plates were incubated at 30°C for 2 days unless noted.

We also tested the effect of tryptophan prototrophy on cells compromised with heat stress. This assay showed that both W303 and BY4741 cells grow well when incubated at 37°C (Fig 4C, left). W303 cell growth was inhibited at 39°C and like CFW and CR treatment; inhibition is completely independent of tryptophan prototrophy (Fig 4C, right). BY4741 cells, however, do not show growth sensitivity at 39°C (Fig 4C, right, row 7).

In converse to CFW, CR and heat stress, the presence of wild type *TRP1* was able to recover cell growth sensitivity due to TTO in prototrophic cells. W303 prototrophic cells were able to overcome growth inhibition due to 0.15% TTO if they contain *his3-11,15*, but not *trp1-1* (Fig 4D, row 5 and 6). The W303 cells containing several auxotrophic markers could not grow in the presence of 0.15% TTO, nor could the BY4741 cells indicating that multiple auxotrophies are also detrimental for the TTO response (Fig 4D, rows 7 and 8). While the effects of TTO are not the same as for SDS, they indicate that tryptophan prototrophy has a similar trend on growth recovery to cells compromised with TTO as with SDS. These results indicate that the activity of the CWI pathway is independent of tryptophan synthesis. Perhaps the stress response involving tryptophan prototrophy is particular to membrane disruptions as opposed to cell wall perturbations.

The *TRP1* gene product is essential for yeast cells to biosynthesize tryptophan. *S. cerevisiae* uses a shared pathway to synthesize tryptophan that also synthesizes phenylalanine and tyrosine (Fig 5A). We know that cells mutant for any of the enzymes specific to the tryptophan biosynthesis pathway are sensitive to SDS (33). We wanted to know if either phenylalanine or tyrosine auxotrophy causes the same response to SDS as an aberrant *TRP1* gene.

**Fig 5.**
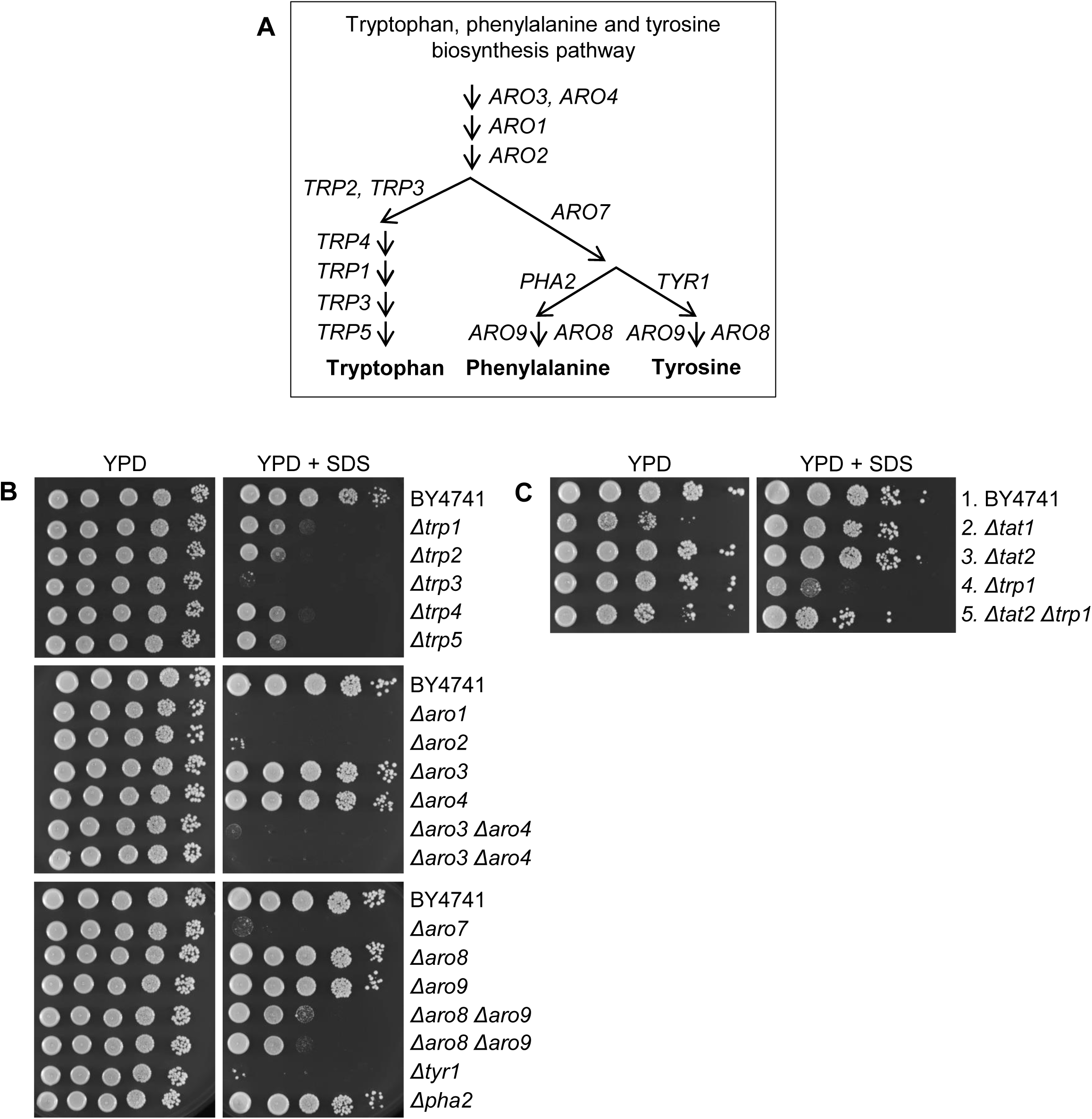
Cells deficient in the biosynthesis of tryptophan or tyrosine are sensitive to SDS. (A) Schematic of the tryptophan, phenylalanine and tyrosine biosynthesis pathway. (B) BY4741 cells (*hisΔ1 leu2Δ0 met15Δ0 ura3Δ0)* and cells harboring specified deletions in the tryptophan, phenylalanine and tyrosine biosynthesis pathway were 10-fold serially diluted onto YPD or YPD containing 0.0075% SDS and incubated at 30°C for 2 days. (C) The indicated BY4741 cells were assayed the same as in B.

To examine this further we obtained BY4741 haploid yeast deletion cells from the EUROpean Saccharomyces Cerevisiae Archive for Functional Analysis (EUROSCARF) (46) harboring a gene deletion for one of each of the enzymes involved in the tryptophan, phenylalanine and tyrosine biosynthesis pathway. From this set of mutants, we also created *Δaro3 Δaro4* and *Δaro8 Δaro9* double deletion mutants. A serial dilution assay shows that growth of any of the cells defective in tryptophan, phenylalanine or tyrosine biosynthesis grow healthy and robust on YPD plates while their growth varies with the addition of SDS (Fig 5B). As expected, all of the mutants specific to the tryptophan branch, *Δtrp1, Δtrp2, Δtrp3, Δtrp4* or *Δtrp5*, grew poorly when compromised with SDS where growth of *Δtrp3* cells were the most inhibited. *ARO1, ARO2, ARO3* and *ARO4* encode for enzymes that are the common precursor to all three amino acids. Growth of the *Δaro1* or *Δaro2* cells was severely debilitated by SDS. *ARO3* and *ARO4* encode enzymes that can substitute for each other and as predicted, the *Δaro3* or *Δaro4* cells show robust growth whereas growth of the *Δaro3 Δaro4* double deletion was extremely impaired by SDS. Growth of *Δaro7*, the common precursor to the phenylalanine and tyrosine branch, was almost completely arrested by SDS. SDS also acutely inhibits the mutant cells specific to the tyrosine branch, *Δtyr1*. This is in contrast to the cells specific to the phenylalanine branch, *Δpha2*, which showed robust growth in the presence of SDS. *ARO8* and *ARO9* encode enzymes that catalyze the last step in the synthesis of tyrosine and phenylalanine. These two enzymes are also able to catabolize tryptophan, tyrosine or phenylalanine and make precursors utilized by Tyr1p and Pha2p. We show that SDS did not compromise growth of *Δaro8* or *Δaro9* single mutant cells whereas the double mutant cells, *Δaro8 Δaro9*, were mildly inhibited by the same treatment.

This experiment uncovers that tyrosine is also required for yeast cells to survive SDS-treatment along with a fully functional tryptophan biosynthesis pathway and provides additional evidence that growth is influenced by tryptophan biosynthesis enzymes.

It has been shown before that tryptophan can be imported through channels other than Tat2p, primarily through Gap1p (18). We retrieved *Δtat2* cells from the EUROSCARF deletion library and compared their growth to *Δtrp1* cells in the presence of SDS. In contrast to *Δtrp1*, *Δtat2* cell growth was not inhibited by SDS treatment (Fig 5C, row 3). We constructed a double deletion mutant, *Δtat2 Δtrp1*, which would not be able to import tryptophan through Tat2p nor make its own. Growth of the *Δtat2 Δtrp1* double deletion mutant was mildly inhibited on SDS containing plates compared to the single mutant, *Δtat2.* Yet, compared to the *Δtrp1* single mutant, the additional loss of *Δtat2* conferred some growth resistance towards SDS treatment indicating that indeed tryptophan uptake is maintained without Tat2p (Fig 5C, rows 3, 4 and 5). In this same assay, we tested the growth of *Δtat1* cells (Fig 5C, row 2). *Tat1* encodes for the high affinity tyrosine permease. We found that *Δtat1* cell growth was not affected by SDS suggesting that tyrosine uptake is also remediated by different means during an SDS response. These results suggest that internal tryptophan and tyrosine levels are important during an SDS assault and that they are acquired through uptake systems other than Tat2p and Tat1p. Because *Δtrp1* cell growth is so constrained by SDS suggest that utilization of the tryptophan biosynthesis pathway is also significant during an SDS response.

### SDS treatment enhances tryptophan uptake

We considered that membrane disruptions caused by SDS could interrupt tryptophan uptake systems and this is why *TRP1* prototrophy is imperative to cells compromised with SDS. If SDS inhibits amino acid import, our results indicate that it is specific for tryptophan and also for tyrosine. To determine tryptophan uptake, we used prototrophic W303 cells whose growth is uncompromised on YPD plates containing 0.0075% SDS (Fig 4A, row 4). We found that import of radiolabeled L-[5-3H]tryptophan or L-[2,5-3H]histidine was enhanced within minutes upon 0.0075% SDS exposure compared to uncompromised cells in liquid culture (Fig 6). It is possible that tryptophan and histidine leak into cells through membrane holes created by SDS at this concentration and that is the explanation for enhanced uptake. However, we found the same enhanced uptake when we challenged cells with a lower concentration of SDS at (0.005%. These results suggest that cells are not starved for tryptophan or histidine upon SDS administration. It has been shown that Gap1p activity can be produced within 5 minutes under certain conditions (30-32). It is possible that Gap1p as a high capacity permease is activated by SDS treatment. Because tryptophan uptake is not inhibited by SDS treatment, provides further evidence that need for tryptophan itself is important.

**Fig 6.**
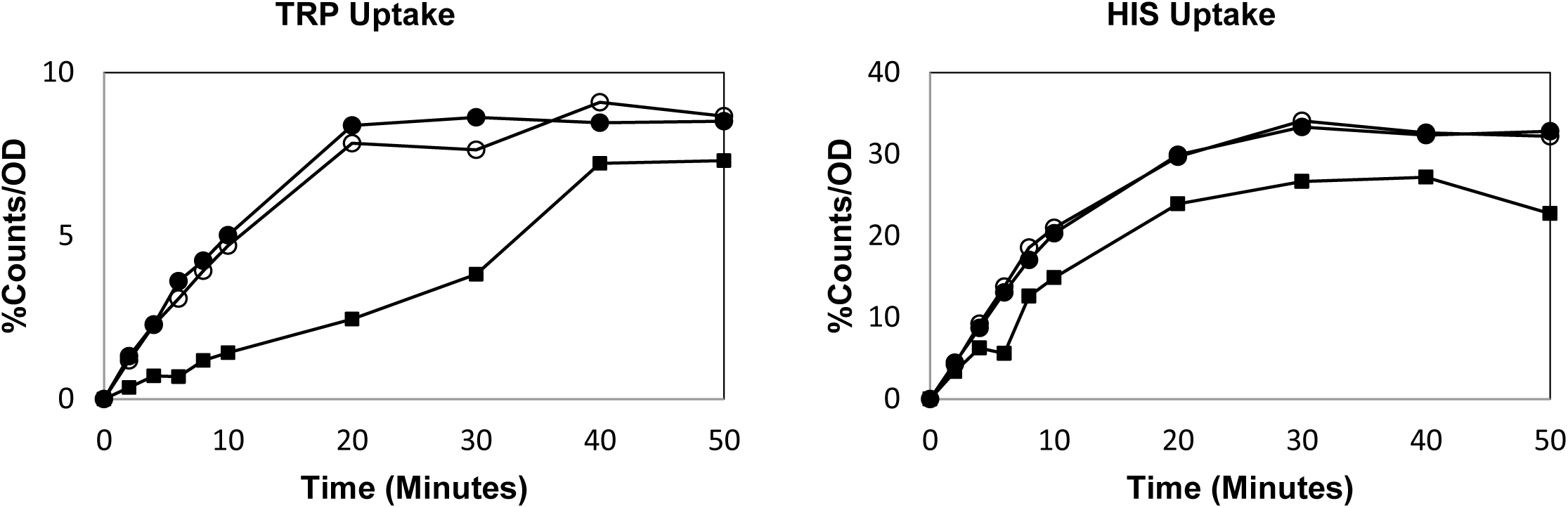
SDS enhances tryptophan uptake. The import rates of radiolabeled tryptophan and histidine were measured using W303 prototrophic cells (*ADE2, HIS3, LEU2, TRP1, and URA3)* in the absence (squares) or presence of (open circles) 0.0075% SDS or (solid circles) 0.005% SDS (See materials and methods). Shown are representative curves.

Because we recovered the *TAT2* tryptophan permease from a suppressor gene screen using *Δmck1* cells in the presence of SDS, we studied tryptophan in the recovery from cell membrane stress due to SDS exposure. First, we show that cells harboring *Δtrp1* have a clear disadvantage in the response to SDS compared to auxotrophies for adenine, histidine, leucine or uracil. Next, we found that tryptophan prototrophy is also critical for stress tolerance towards TTO, another membrane destabilizing drug. While both SDS and TTO cause CWI activation, we demonstrate that tryptophan prototrophy is not able to alleviate growth inhibition due to other cell wall/membrane damaging treatments that also activate the pathway indicating a distinction from CWI signaling. This also implicates that the resistance to growth inhibition shown by tryptophan prototrophic cells may be specific to the type of membrane damage created by SDS and TTO as opposed to cell wall disruptions. In addition, we uncover that tyrosine biosynthesis is also important for resistance to SDS-induced growth inhibition whereas phenylalanine biosynthesis is dispensable. We also found that in the presence of SDS, *Δtat2* deletion cells show increased growth resistance to *Δtrp1* cells indicating that internal tryptophan levels are maintained during an SDS assault through uptake systems other than Tat2p. Finally, we observe that both tryptophan and histidine import becomes enhanced immediately upon addition of SDS as a further indication that SDS-induced growth inhibition is not due to nutrient starvation in general.

These results suggest that tryptophan, tryptophan biosynthesis and tyrosine biosynthesis play a role in the plasma membrane stress response. It is thought that the constitutive permeases, such as Tat2p and Tat1p, uptake amino acids for use in protein synthesis. Gap1p, however, is a transporter of all amino acids and is regulated by nitrogen (47) therefore it is thought that Gap1p acquires amino acids for use as a nitrogen source (48). Future directions would be to explore tryptophan and tyrosine biosynthesis and catabolism in respect to nitrogen response pathways and cell membrane damage.

## Materials and methods

### Construction of yeast strains

The yeast cells used are derivatives of W303 (strain list in Table 1) except for the BY4741 control cells or BY4742 Mat α derivative from this laboratory and the haploid BY4741 deletion cells from EUROSCARF (46). The LSY112, LSY113, LSY114, LSY115, LSY116, LSY118, LSY119, LSY121 and LSY123 cells were made by standard cross and dissection procedures using RUY508 Mat <u>a</u> *his3-11,15 leu2-3,112 trp1-1 ura3-1 can1-100* (this laboratory) crossed with prototrophic parent cells, Mat α *ade2-1 can1-100*, which only contain the indicated selectable markers. The haploid genotypes were determined by tetrad analysis. The prototrophic parent cells, kindly provided by Fred Cross, were made in W303 by transformation with the various cloned markers (*ADE2, HIS3, LEU2, TRP1*, and *URA3*) and using mating and tetrad analysis to get haploids with the different combinations. The double mutants in Fig 5, LSY203, LSY204 (*Δaro3 Δaro4)* and LSY208, LSY209 (*Δaro8 Δaro9)* which are independent clones and LSY200 (*Δtat2 Δtrp1)* were made using the EUROSCARF haploid BY4741 deletion cells (*Δaro4*, *Δaro8* or *Δtat2*) crossed with BY4742 to obtain Mat α cells containing the desired deletion and then crossed with the respective secondary deletion cells from the EURPSCARF BY4741 library (*Δaro3, Δaro9* or *Δtrp1*). The double mutant genotypes were determined with tetrad analysis and PCR. The *TAT2*/pRS425 plasmid was made by PCR of BG1805 YORF (YOLO20W) from the start to the stop codon of *TAT2* using the primers FP1 (CCTGCAGCCCGGGGGATCCA) and RP1 (GGCGGCCGCTCTAGAACTAGTTAACACCAGAAATGGAACT). The *TAT2* PCR product was assembled into the pRS425-2micron *LEU* marked plasmid by Gibson Assembly using the universal primers for pRS425, FP2 (CTAGTTCTAGAGCGGCCGCC) and RP2 (TGGATCCCCCGGGCTGCAGG). The construction of BCY061 (*mck1::KanMX)* has been described in (12). The *TAT2*/pRS425 plasmid or empty vector was transformed into BCY061) using standard lithium acetate transformation and selected on synthetic glucose medium lacking leucine.

**Table 1.** S. cerevisiae strains used in this study

| **Strain** | **Genotype** | **Source** |
| --- | --- | --- |
| RUY508 | MAT<sub>a</sub> *his3-11,15 leu2-3,112 trp1-1 ura3-1 can1-100* | This Laboratory |
| BCY061 | MAT<sub>a</sub> *mck1::KanMX his3-11,15 leu2-3,112 trp1-1 ura3-1 can1-100* | (12) |
| LSY183 | MAT<sub>a</sub> *trp1::KanMX hisΔ1 leu2Δ0 met15Δ0 ura3Δ0* | Euroscarf |
| RUY120 | MAT<sub>a</sub> *ade2-1 his3-11,15 leu2-3,112 trp1-1 ura3-1 can1-100* | This Laboratory |
| RUY064 | MAT<sub>a</sub> *hisΔ1 leu2Δ0 met15Δ0 ura3Δ0* | This Laboratory |
| RUY065 | MAT<sub>alpha</sub> *hisΔ1 leu2Δ0 met15Δ0 ura3Δ0* | This Laboratory |
| LSY119 | MAT<sub>a</sub> *can1-100* | This Laboratory |
| LSY123 | MAT<sub>a</sub> *trp1-1 can1-100* | This Laboratory |
| LSY116 | MAT<sub>a</sub> *ade2-1 his3-11,15 leu2-3,112 ura3-1 can1-100* | This Laboratory |
| LSY112 | MAT<sub>a</sub> *ade2-1 his3-11,15 leu2-3,112 trp1-1 ura3-1 can1-100* | This Laboratory |
| LSY113 | MAT<sub>a</sub> *his3-11,15 leu2-3,112 trp1-1 ura3-1 can1-100* | This Laboratory |
| LSY114 | MAT<sub>a</sub> *ade2-1 leu2-3,112, trp1-1 ura3-1 can1-100* | This Laboratory |
| LSY115 | MAT<sub>a</sub> *ade2-1 his3-11,15 trp1-1 ura3-1 can1-100* | This Laboratory |
| LSY118 | MAT<sub>a</sub> *ade2-1 his3-11,15 leu2-3,112 trp1-1 can1-100* | This Laboratory |
| LSY121 | MAT<sub>a</sub> *his3-11,15 can1-100* | This Laboratory |
| LSY176 | MAT<sub>a</sub> *aro1::KanMX hisΔ1 leu2Δ0 met15Δ0 ura3Δ0* | Euroscarf |
| LSY177 | MAT<sub>a</sub> *aro2::KanMX hisΔ1 leu2Δ0 met15Δ0 ura3Δ0* | Euroscarf |
| LSY178 | MAT<sub>a</sub> *aro3::KanMX hisΔ1 leu2Δ0 met15Δ0 ura3Δ0* | Euroscarf |
| LSY179 | MAT<sub>a</sub> *aro4::KanMX hisΔ1 leu2Δ0 met15Δ0 ura3Δ0* | Euroscarf |
| LSY203 | MAT<sub>a</sub> *aro3::KanMX aro4::KanMX hisΔ1 leu2Δ0 met15Δ0 ura3Δ0* | This Laboratory |
| LSY204 | MAT<sub>a</sub> *aro3::KanMX aro4::KanMX hisΔ1 leu2Δ0 met15Δ0 ura3Δ0* | This Laboratory |
| LSY180 | MAT<sub>a</sub> *aro7::KanMX hisΔ1 leu2Δ0 met15Δ0 ura3Δ0* | Euroscarf |
| LSY181 | MAT<sub>a</sub> *aro8::KanMX hisΔ1 leu2Δ0 met15Δ0 ura3Δ0* | Euroscarf |
| LSY182 | MAT<sub>a</sub> *aro9::KanMX hisΔ1 leu2Δ0 met15Δ0 ura3Δ0* | Euroscarf |
| LSY208 | MAT<sub>a</sub> *aro8::KanMX aro9::KanMX hisΔ1 leu2Δ0 met15Δ0 ura3Δ0* | This Laboratory |
| LSY209 | MAT<sub>a</sub> *aro8::KanMX aro9::KanMX hisΔ1 leu2Δ0 met15Δ0 ura3Δ0* | This Laboratory |
| LSY183 | MAT<sub>a</sub> *trp1::KanMX hisΔ1 leu2Δ0 met15Δ0 ura3Δ0* | Euroscarf |
| LSY184 | MAT<sub>a</sub> *trp2::KanMX hisΔ1 leu2Δ0 met15Δ0 ura3Δ0* | Euroscarf |
| LSY185 | MAT<sub>a</sub> *trp3::KanMX hisΔ1 leu2Δ0 met15Δ0 ura3Δ0* | Euroscarf |
| LSY186 | MAT<sub>a</sub> *trp4::KanMX hisΔ1 leu2Δ0 met15Δ0 ura3Δ0* | Euroscarf |
| LSY187 | MAT<sub>a</sub> *trp5::KanMX hisΔ1 leu2Δ0 met15Δ0 ura3Δ0* | Euroscarf |
| LSY188 | MAT<sub>a</sub> *tyr1::KanMX hisΔ1 leu2Δ0 met15Δ0 ura3Δ0* | Euroscarf |
| LSY189 | MAT<sub>a</sub> *pha2::KanMX hisΔ1 leu2Δ0 met15Δ0 ura3Δ0* | Euroscarf |
| LSY200 | MAT<sub>a</sub> *tat2::KanMX trp1::KanMX hisΔ1 leu2Δ0 met15Δ0 ura3Δ0* | This Laboratory |

### Cell culture and media

Yeast cultures were grown in standard complete Yeast extract and Peptone medium containing 2% Dextrose (YPD). Plates were made with 25mL of YPD and 2% agar. The chemicals were added to YPD plates with final the concentration of 0.0075% SDS, 10ug/ml Calcofluor White (Sigma 18909), or 10ug/ml Congo Red, (Sigma C-6767) or 0.15% Tea Tree Oil (Sigma SMB00386). 0.5% Tween 40 was added to the Tea Tree Oil and control plates to assure solubility of the Tea Tree Oil. In Fig 1B, supplements were spread on top of YPD plates at 1X final concentration from 100X stocks supplied at 6g/L adenine, 2g/L histidine, 12g/L leucine, 8g/L tryptophan or 2g/L uracil. In Fig 2C, the supplements were spread on top of YPD plates with 1X, 2X, 4X increments of 100X tryptophan at 8g/L or 100X histidine at 2g/L. All plates were incubated at 30°C for two days unless otherwise indicated. Images are representative of three independent experiments.

### Suppressor gene screening

The suppressor gene screen, which has been described in (49), was done using *mck1*::KanMX BCY061 cells transformed with the ATCC YEp13 total yeast genomic DNA library cloned in a 2-micron/*LEU2* high-copy plasmid (50).

### Serial dilution

Cells were grown in YPD at room temperature overnight. Cell concentrations were normalized by OD595, diluted serially 10-fold for five dilutions before plating. Plates were incubated at indicated temperatures for 2 days.

### Amino acid uptake assay

The protocol was adapted from J Heitman (51). LSY119 cells in log-phase were harvested and washed once with 10mM sodium citrate, pH4.5, and resuspended in 50mLs of 10mM sodium citrate, pH4.5, containing 20mM ammonium sulfate and 2% glucose. SDS at 0.0075% was added and the 0 time point was taken immediately before the radioactive substrate addition. Uptake was assayed by adding 0.5mL of radiolabeled amino acid mixture to 4.5mL cell culture. The radiolabeled amino acid mixture was made with 495.5uL H2O and 4.5uL of either tryptophan, L-[5-3H], 20 Ci/mmol, or histidine, L-[ring-2,5-3H], 50 Ci/mmol (American Radiolabeled Chemicals, Inc.). Aliquots of 0.5mL were taken at 0 time point (no radioactivity) and in 2 min intervals from 2-10 min and then every 10 min up to 50 min. Cells were vacuum filtered onto Whatman glass microfiber filters (Sigma WHA1825025) presoaked in 10mM sodium citrate, pH4.5, and washed twice with 2mL10mM sodium citrate, pH4.5, containing 2mM tryptophan and histidine. Filters were dried and the remaining radioactivity was quantified with 5mL scintillation fluid. Percent counts were normalized by cell density determined by OD_595_.

### Reagent availability

All strains and protocols are available upon request.

## Acknowledgement

We thank Dr. Frederick Cross for providing us yeast strains.

